# Metabolomic Selection for Enhanced Fruit Flavor

**DOI:** 10.1101/2020.09.17.302802

**Authors:** Vincent Colantonio, Luis Felipe V. Ferrão, Denise Tieman, Nikolay Bliznyuk, Charles Sims, Harry Klee, Patricio R. Munoz, Marcio F. R. Resende

**Author notes:** Contributed equally (Vincent Colantonio, Luis Felipe V. Ferrão).

## Abstract

Although they are staple foods in cuisines globally, commercial fruit varieties have become progressively less flavorful over time. Due to the cost and difficulty associated with flavor phenotyping, many breeding programs have long been challenged in selecting for this complex trait. To address this issue, we leveraged targeted metabolomics of diverse tomato and blueberry accessions and their corresponding consumer panel ratings to create statistical and machine learning models that can predict sensory perceptions of fruit flavor. Using these models, a breeding program can assess flavor ratings for a large number of varieties, previously limited by the low-throughput and high cost of consumer sensory panels. The ability to predict consumer ratings of liking, sweet, sour, umami, and flavor intensity was evaluated by a 10-fold cross-validation and the accuracies of 18 different models are assessed. The best performing models were used to infer the flavor compounds (sugars, acids, and volatiles) that contribute most to each flavor attribute. The prediction accuracies were high for most attributes in both blueberries and tomatoes. We expect that these models will enable an earlier incorporation of flavor as breeding targets and encourage selection and release of more flavorful fruit varieties.

## 1 Introduction

Plant breeders and geneticists have made continuous and substantial progress in the development of varieties that are more resilient and higher yielding - much to the benefit of producers worldwide. Yet, during this extended period of progress, consumer-oriented quality traits such as flavor have been neglected or treated as low priority breeding targets, contributing to widespread consumer dissatisfaction with modern fruit and vegetable varieties (1). An important reason for this low priority is a reward system that pays growers based on crop yield, leading to prioritization of breeding targets that are mainly producer oriented. However, as consumer willingness to pay premiums for higher quality products rises, demand for consumer-oriented traits in food production systems is increasing (2). Accordingly, a reemerging interest in fruit and vegetable flavor quality creates the need for high-yielding varieties with exceptional flavor profiles.

Fruit flavor is the product of the complex interactions between the chemical composition of a fruit, and the taste, olfaction, and psychology of the consumer (3; 4; 5). To breed and develop varieties with improved flavor properties, the genetic complexities of fruit flavor must be captured and assessed. Flavor is currently evaluated by consumer sensory panels or individually by breeders. Field evaluations are generally subjective and error-prone as they typically consist of the sensory preferences of one or few individuals. However, field evaluation has an advantage in that many varieties can be evaluated in a given day. In contrast, sensory panels are more objective, accurate, and well established, but they can be costly, time consuming, and difficult to scale to a large breeding program. The difficulties associated with accurate flavor phenotyping have contributed to the lack of selection for fruit flavor, and thereby contributed to the widespread consumer belief that commercial fruit flavor has declined (6; 7). Cheap and scalable flavor selection methods would greatly benefit the breeding process.

The main driver of fruit flavor perception is its chemical composition. Fruits contain a diverse array of sugars, acids, and volatiles whose concentrations are driven by genetic and environmental effects. Sugars and acids are largely perceived by taste receptors on the tongue, and the volatiles by receptors located in the olfactory epithelium (8). We hypothesized that by quantifying the chemical profile of a fruit and its corresponding consumer perception, models predicting consumer flavor preferences can be created. By eliminating the cost associated with consumer sensory panels, these prediction models can increase throughput of flavor phenotyping, allowing a breeder to make selections for improved flavor on hundreds of varieties per season. Leveraging the trained models for inference, specific metabolites that underlie consumer flavor preferences can also be elucidated, identifying targets for breeding and the food industry to enhance flavor in their products. This approach is analogous to the concept of genomic selection (9), where DNA markers are used in plant breeding programs to predict highly complex traits. Here, we propose the use of statistical methods to model the metabolomic profile in a breeding population and predict flavor perception.

In flavor studies, the most widespread statistical modeling approaches to date include multiple linear regression and partial least squares regression (10). However, the process of developing metabolomic-based prediction models can be challenging due to the large number of chemical compounds present in a fruit, and the fact that the concentrations of many of the flavor-associated chemicals are correlated with one another due to shared biosynthetic pathways. Fortunately, breeders and quantitative geneticists are already dealing with similar types of data in the area of genomic selection (GS) to make selection of complex traits using genomic information. Thus, there are several statistical and machine learning prediction models that have been tested in GS and other prediction areas. Based on the model assumptions and specifications, different prediction accuracies and abilities to infer the contributing features are expected. With the advent of genomic selection, for example, a variety of Bayesian linear regression models with different priors were proposed to predict complex phenotypes using DNA markers. These models included Bayes A and Bayes B (9), Bayes Cπ (11), Bayesian LASSO (12), and Bayesian Ridge Regression (13; 14). Recently, there has been increasing interest in machine learning models for genomic and bioinformatic applications. However, few studies have applied these models at the metabolome level, or specifically for the enhancement of fruit flavor.

Here we address the limitations in flavor phenotyping and propose a phenotyping approach that has significantly higher throughput compared to current standards. We assessed a range of statistical and machine learning models that take the chemotype of a fruit and make predictions of its consumer flavor perception. To this end, we combined information at the metabolome and sensory panel level for two important horticultural crops, tomato and blueberry, and demonstrate that metabolomic prediction models can be employed in a breeding program to make simpler and more accurate selections for flavorful varieties. Additionally, we leverage the trained models to infer the contributions of volatiles, sugars and acids to sensory perceptions and consumer likeability. Our results suggest that up to 55% of the variance associated with overall consumer liking can be attributed to volatile compounds. Furthermore, we demonstrate that machine learning approaches are generally the best predictors of consumer flavor preferences, and metabolomic selection accuracies are superior to genomic selection models, highlighting the potential in breeding applications.

## 2 Results

### 2.1 The data

In order to study the capacity of different prediction models and the importance of different metabolites in flavor perception, we performed an analysis of previously published data (15; 16; 4; 5) with new data added in this study, for two fruit species: tomato (*Solanum lycopersicum*) and blueberry (*Vaccinium* spp.).

For each fruit, targeted sets of sugars, acids, and volatiles were quantified in diverse accessions including commercial cultivars, heirloom varieties, and germplasm selections from the University of Florida tomato and blueberry breeding programs. For tomato, 68 sugars, acids, and volatiles were analyzed in 147 genotypes grown and evaluated in multiple seasons, for a total of 209 samples. For blueberry, 63 accessions evaluated in multiple seasons for a total of 244 samples, were analyzed for firmness and 55 sugars, acids, and volatiles. In addition, consumer sensory panels rated each accession for flavor attributes including sweetness, sourness, flavor intensity, and overall liking. Additionally, sensory perceptions of umami were quantified solely for tomato.

### 2.2 Network analysis recapitulates metabolic pathways

A weighted correlation network analysis was performed on the metabolite contents across all fruit accessions for tomato and blueberry (Figure 1A and Figure 2A). The results are largely consistent with knowledge of the individual biosynthetic pathways and provide new insights into the relationships between pathways. For example, there are strong associations between the apocarotenoid volatiles (e.g., geranial and *β*-cyclocitral) and the fatty acid-derived volatiles (e.g., 1-pentanol and *E*-2-heptenal). Apocarotenoid volatiles are derived from precursors localized in plastids and their contents substantially increase during the conversion of chloroplasts to chromoplasts (17), while the precursors of fatty-acid-derived volatiles are membrane lipids. *TOMLoxC* is an essential enzyme for the synthesis of five and six carbon fatty acid-derived volatiles (18; 19). Recently, a potential link between these pathways was recently proposed with a QTL analysis that implicated a role for *TOMLoxC* in apocarotenoid synthesis possibly by co-oxidation mechanism (20).

**Figure 1:**
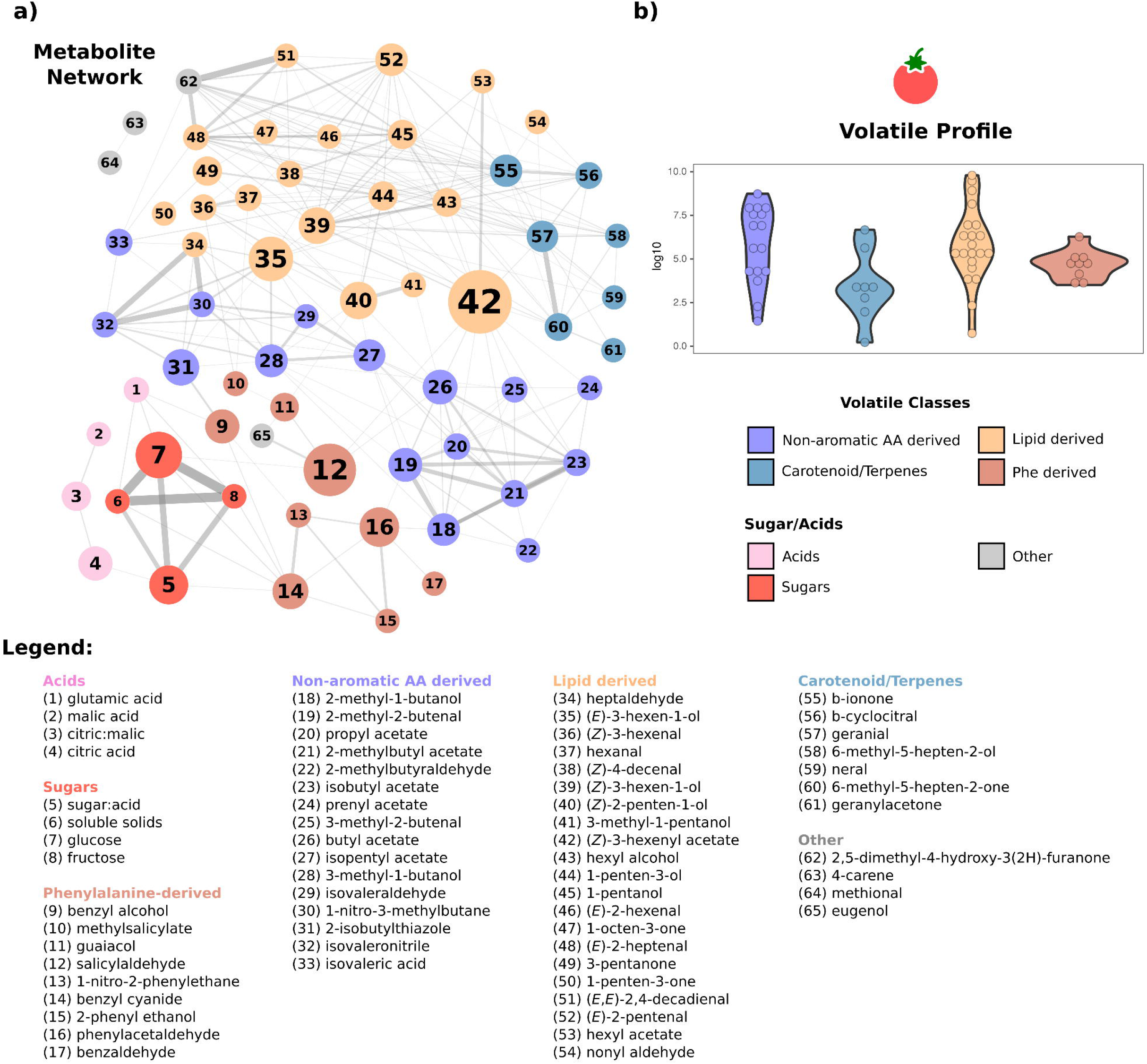
(a) Weighted correlation network analysis of tomato metabolites and their assigned clusters based on their known biochemical classification. The size of each metabolite node indicates betweenness centrality. The size of each edge indicates correlation. The identity of each metabolite is denoted by number in the legend. (b) Distribution of metabolite concentrations for each group across tomato accessions.

### 2.3 Contributions of sugars, acids, and volatiles to flavor perceptions

In order to determine if the fruit metabolome could explain variation in sensory panel ratings, we first partitioned the metabolome into two main groups: sugars/acids and volatiles. We separated the consumer sensory variance into aggregated components explained by each group. In both tomato and blueberry, a large proportion of the variance was explained by sugars/acids and volatiles, while little variance was attributed to the residuals (Figure 3). These results suggest that the fruit metabolome is predictive of corresponding consumer taste panel ratings. Furthermore, the proportion of variance explained by sugars/acids varied across the flavor attributes and contrasted between the two species. For instance, 77% of the tomato sourness variance was explained by the content of sugars/acids while these compounds only explained 42% of blueberry sourness. Similarly, while sugars/acids predominantly (60%) explained blueberry sweetness, a larger portion of the tomato phenotypic variance (59%) could be explained by the volatiles. As previously described (3), the results indicate the large influence that volatile compounds can have on sensory attributes in both species, which in turn, highlights how important these compounds are to breeding programs for improvement of fruit flavor. For example, the variance decomposition of overall liking score estimates that 41% and 55% of the variance was explained by volatile organic compounds in tomato and blueberry, respectively.

To further understand how the fruit chemical profile affected consumer flavor, we partitioned the metabolites into modules according to their biochemical classifications (Figures 1, 2; Supplemental Tables 1 and 2).A linear mixed model was fitted considering each module as a random effect and their corresponding proportions of variance were obtained (Figure 3; Supplemental Figure 1). The sugar module was a strong driver of liking (43% in tomato and 19% in blueberry) and sweetness (32% in tomato and 27% in blueberry), while the module representing acids drove sourness (49% in tomato and 38% in blueberry) in both fruits. Some volatile modules were found to make large contributions to flavor ratings. For instance, phenylalanine-derived and lipid-derived compounds contributed to sweetness perception (25 and 23% respectively) and overall liking score (15 and 12% respectively) in tomato. Lipid derived volatiles and compounds grouped as carotenoid/terpenes explained 15 and 21% of blueberry overall liking score, respectively. These results are consistent with previous results that showed strong positive correlations of specific volatiles with fruit sweetness (16; 4).

**Figure 2:**
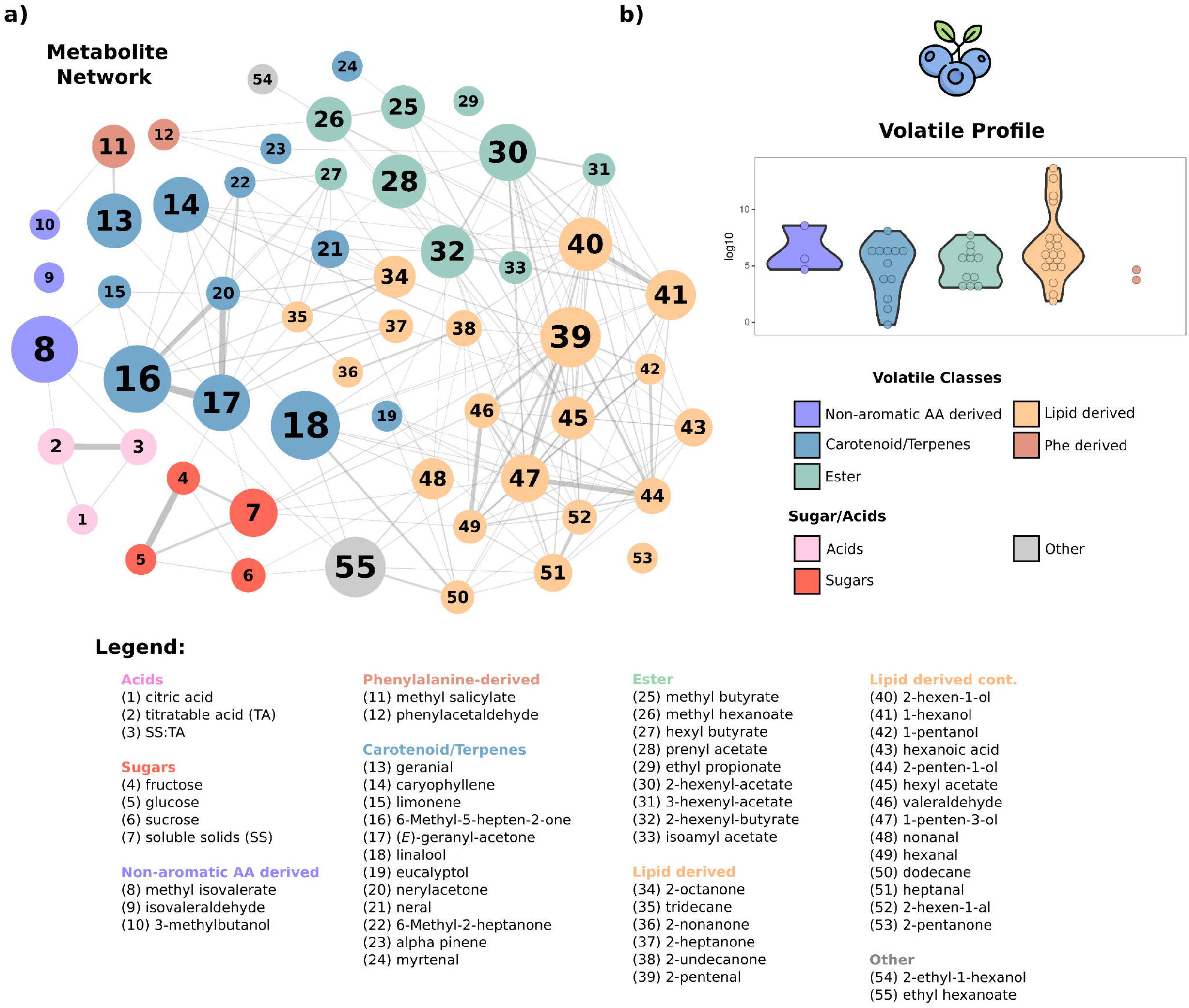
(a) Weighted correlation network analysis of blueberry metabolites and their assigned clusters based on their known biochemical classification. The size of each metabolite node indicates betweenness centrality. The size of each edge indicates correlation. The identity of each metabolite is denoted by number in the legend. (b) Distribution of metabolite concentrations for each group across blueberry accessions.

**Figure 3:**
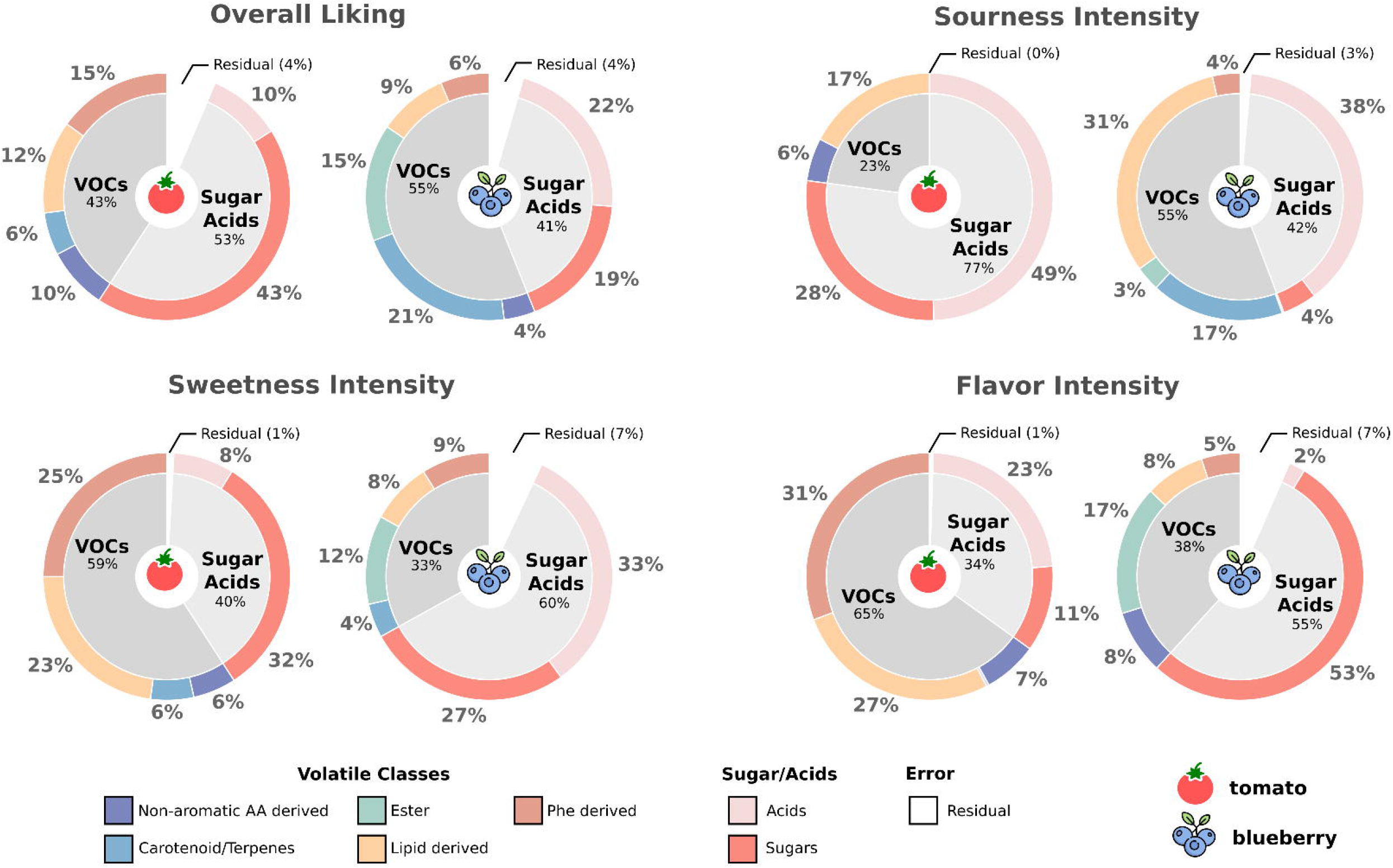
Variation in sensory panel ratings explained by sugars/acids and volatiles overall, and by groups of metabolites of known biochemical classification in tomato and blueberry

### 2.4 Predicting consumer preference

Eighteen statistical and machine learning methods were employed to predict sensory traits from sugar, acid, and volatile concentrations. Each model was evaluated in a 10-fold cross validation and each fold was assessed by the correlation between predicted and observed consumer taste panel ratings (Figure 4A, Supplemental Figure 1). The cross validation was repeated 10 times and results were averaged for the final prediction accuracies. We observed the highest prediction accuracies from the XGBoost, gradient boosting machines, and neural network models. The XGBoost model showed an average improvement of 19% over the linear regression and 11% over partial least squares, models traditionally used in food science applications. The accuracy for the best model (XGBoost) ranged from 0.62 to 0.87 across all traits and in both species. We found the most predictable traits in tomato to be sweetness (0.8), flavor intensity (0.78), and sourness (0.69), and the most predictable traits in blueberry to be sourness (0.87) and sweetness (0.75).

**Figure 4:**
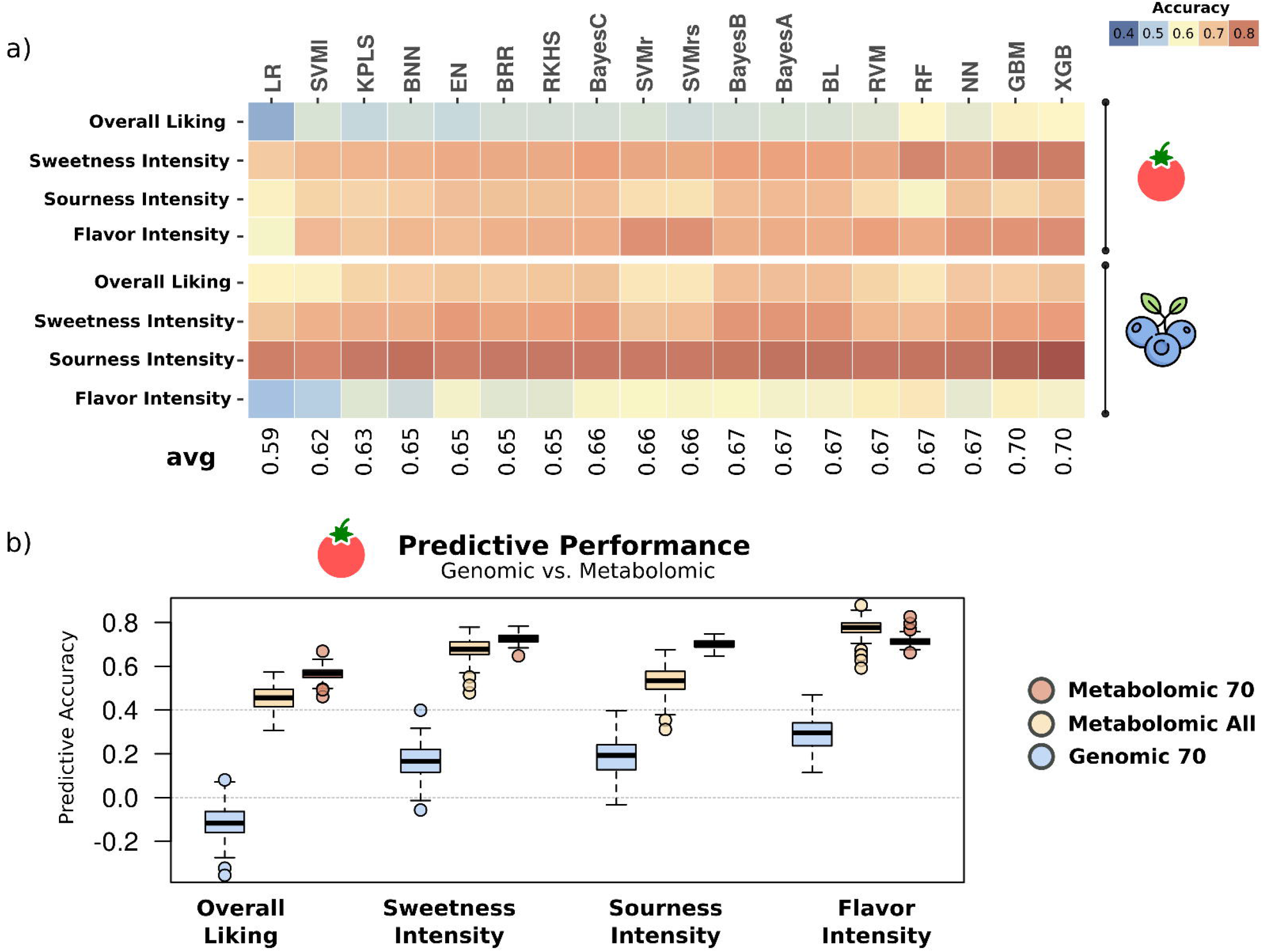
(a) Accuracy of predicting flavor ratings from metabolome data across the follow range of statistical and machine learning models for tomato and blueberry: linear regression(LR), Linear Support Vector Machine (SVMl), Kernel Partial Least Squares (KPLS), Bayesian Neural Net (BNN), Elastic Net (EN), Bayesian Rigge Regression (BRR), BayesA, BayesB, BayesC, Bayesian Lasso (BL), Reproducing Kernel Hilbert Space (RKHS), Support Vector Machine Radial (SVMr), Support Vector Machine Radial Sigma (SVMrs), Relevant Vector Machine (RVM), Random Forest (RF), Neural Network (NN), Gradient Boost Machine (GBM)and X Gradient Boost (XGB). Refer to the “Material and Methods” section for an overview of the methods compared. (b) Accuracy of predicting perception traits for tomato using 70 individuals with genomic (Genomic 70) and metabolomic (Metabolomic 70) data, as well as all 209 individuals with metabolomic data (Metabolomic All).

To further evaluate the opportunity to use metabolomic selection in breeding and to understand its prediction potential compared to traditional genomic selection (GS) methods, we compiled information from 70 varieties of tomato for which we had whole genome sequence, chemical profile, and sensory panel data (5). Using a 10-fold cross validation, we applied the genomic selection GBLUP method (21) to predict the consumer sensory ratings from a subset of 70,000 SNPs (Figure 4B). We then used the metabolomic information for the same 70 varieties and the same cross validation partitioning to predict the panel ratings. These 70 varieties represent a subset of the total 147 accessions. We found that metabolomic selection outperformed genomic selection in the prediction of all these complex traits, especially for sweetness and overall flavor liking. For these traits, the accuracies of metabolomic selection using 70 varieties were 0.68 and 0.45 for sweetness and overall flavor liking, respectively. These traits were poorly predicted by GBLUP with accuracies of 0.18 and −0.11 for sweetness and overall flavor liking, respectively. While these results are not surprising, given that the metabolite data is capturing both genetic and environmental components of fruit flavor, they highlight the complexity of flavor perception as a breeding target, and the potential of metabolomic selection as a phenotyping tool to support breeding programs compared with other available methods such as GS, for example. The accuracies of metabolomic selection in a model trained with 70 genotypes were slightly lower than the accuracies from the full dataset using 209 observations and 147 genotypes. The exception was for flavor intensity where the subset of 70 reached accuracies of 0.77 and was more predictive than the full dataset. This result can be explained by a smaller variety panel that is less variable and easier to be predicted or by the fact that multiple sensory panels were averaged in the subset analysis resulting in a more precise phenotype.

### 2.5 Metabololites associated with desirable flavor

In order to find sugars, acids, and volatiles that enhance or suppress consumer sensory perceptions of flavor, models for each fruit were trained using all samples for which we had metabolome and sensory panel data (209 for tomato and 244 for blueberry). Two contrasting modeling approaches, BayesA and Gradient Boosting Machines, were chosen for further inference analysis. In BayesA, the *β* coefficients indicate the individual additive effect of that chemical free of interactions. This coefficient predicts if a chemical is important for enhancing the flavor attribute (positive value) or decreasing the flavor attribute (negative value). For Gradient Boosting Machines the variable importance represents the marginal effect of that chemical including the interaction effects with other chemicals. Its value was measured by calculating the difference in prediction accuracy between the models trained using all the predictors and models trained with all predictors except one, permuted across all predictors. This value is then scaled between 0 and 100 where 0 is a not an important predictor and 100 is an important predictor. Importance values for Gradient Boosting Machines indicate solely whether a metabolite is important for a given sensory trait while importance values (*β*’s) for Bayes A models indicate whether a metabolite is important for a given trait and if it induces or suppresses the trait responses (Figure 5).

**Figure 5:**
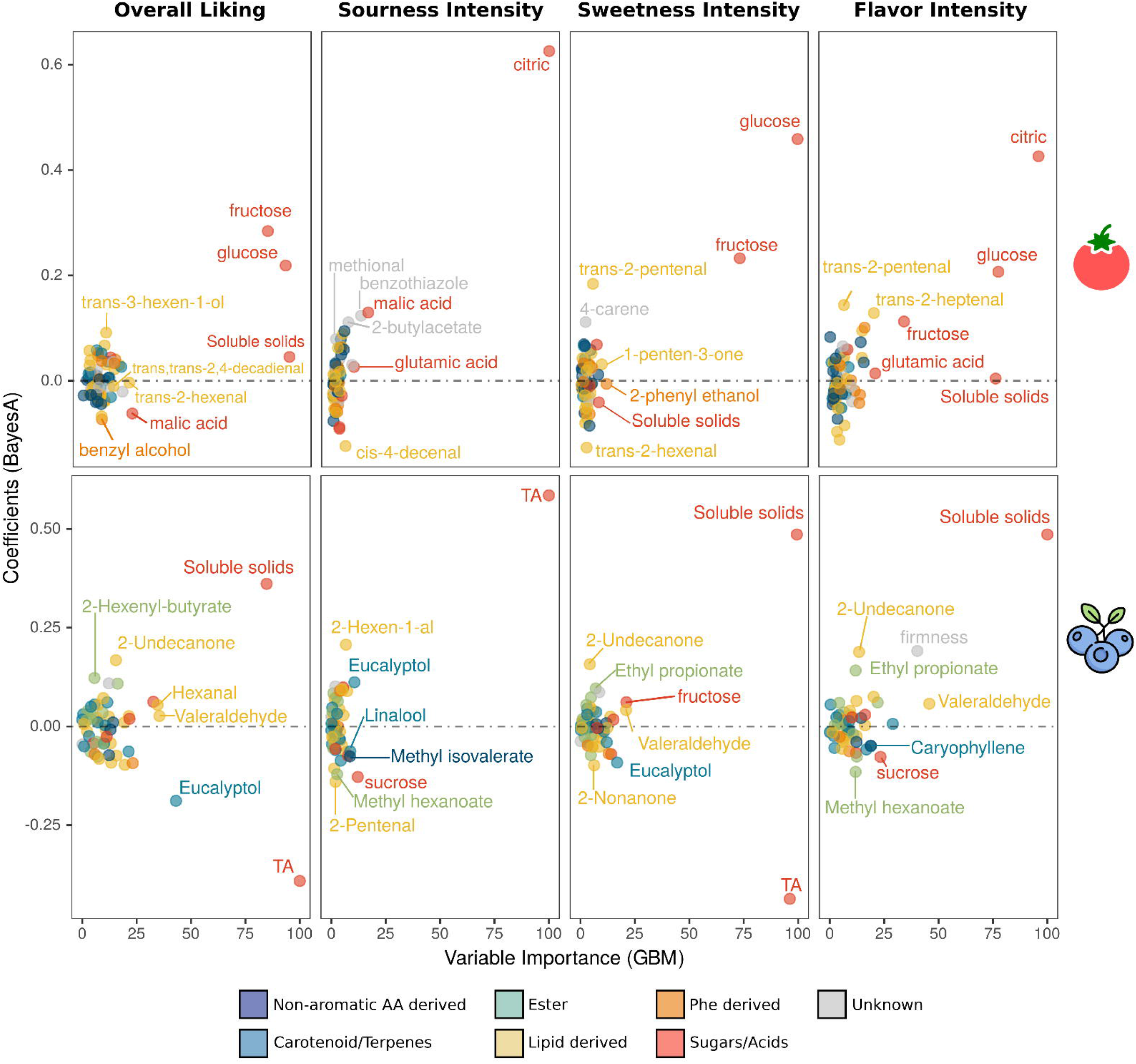
Relative importance estimated with Gradient Boost Machine (x-axis) and effect level estimated with BayesA (y-axis) of each metabolite in predicting ratings of flavor attributes in blueberry and tomato. Colors indicate the type of metabolite. TA, titratable acidity.

For sweetness in tomato, we found glucose and fructose to be the most important sensory perception enhancers. The Gradient Boosting Machines also estimated 1-penten-3-one and 2-phenylethanol to be important for perceived sweetness, while the BayesA model highly ranked two volatiles (*E*-2-pentenal and 4-carene) o be important for sweetness enhancement. *E*-2-pentenal was also found to be an important contributor to overall flavor intensity and umami (Supplemental Figure 1). In blueberry, components important for liking included soluble solids, fructose, and glucose. Additionally, volatiles found to be important for enhancing liking included 2-undecanone, 2-hexenyl-butyrate, and ethyl propionate, while volatiles that were negative to liking included eucalyptol and phenylacetaldehyde. Interestingly, two lipid derived volatiles (2-hexen-1-al and 2-pentenal) had a high positive contribution and the highest negative contributions to sourness in blueberry, respectively. Glutamic acid was highly ranked by both methods as influencing umami perception for tomato, which by definition represents the taste of the amino acid L-glutamate. Two phenylalanine-derived compounds (benzyl cyanide and 2-phenyl ethanol) and one isoleucine/leucine-derived volatile (1-nitro-2-phenylethane) were also highly ranked by Gradient Boosting Machines as umami influencers (Supp. Figure 1). It is important to note, however, that this targeted metabolomic panel is enriched for putative sugar-enhancing compounds and may be limited in the characterization of compounds affecting umami.

## 3 Discussion

Fruit flavor is a complex trait at the intersection between the fruit biochemistry and the consumer sensory perception. Quantification of sensory perception using consumer flavor panels is time and resource consuming and not readily amenable to a high throughput assay, which has hindered plant breeders from selecting for fruit flavor for many years. This has contributed to the widespread decline of consumer satisfaction of many commercial fruit varieties (7; 2). Recently, different high-throughput phenotyping applications were proposed to use 2-D visible light imaging as proxies for plant biomass (Golzarian et al., 2011), reflectance ratios as proxies for yield (22), hyperspectral reflectance as proxies for leaf chlorophyll and nitrogen content (23), and canopy temperature as proxies for drought response (24). Here, and in order to create higher throughput flavor phenotyping methods, we applied statistical and machine learning models that can predict consumer sensory panel ratings from the chemistry of a fruit.

### 3.1 Chemical profiling of a fruit

The cost of targeted and untargeted metabolomics has decreased in recent years (25). In addition, the throughput for metabolite profiling has also increased, which in turn reduces the labor cost per sample. Currently, we are able to analyze 150 samples per day using a purge and trap gas chromatography-mass spectrometry (GC-MS) system. The largest cost in this system is labor to process the fruit and to analyze the data. For the panel generated in this work, we estimate an in-house cost per sample of $10-$20 for GC-MS characterization. This cost allows the characterization of 3 replicates per genotype for costs ranging from $30 - $60 per variety, which is equivalent to the per sample genotyping costs used for genomic selection in many species. If we assume the costs of a flavor sensory panel to be ~ $5000.00 with the evaluation of 5 products ($1000/variety), flavor phenotyping through chemical profiling can be more than 30 times cheaper than sensory evaluations. While these cost estimates assume an in-house analysis and do not consider the capital expenditure to acquire the instrumentation, it highlights the per sample cost reduction over the years and the feasibility of its application in plant breeding programs. Furthermore, with the identification of a set of candidate metabolites that have more influence in a specific flavor attribute of interest, targeted metabolomics could be utilized to further reduce the cost.

### 3.2 Applications of metabolomic selection to breeding of fruits and vegetables

One alternative to phenotype fruits and vegetables for flavor attributes is the establishment of consumer sensory panels. This approach is expensive with low throughput because a sensory panel can usually only taste a limited number of samples (4 to 6) per day. The tomato and blueberry breeding programs at the University of Florida have been using consumer sensory panels to guide breeding decisions for several years (16; 5). To do this, fruit varieties ranging from heirloom and commercial cultivars to varieties currently in development by the breeding programs are subjected to biochemical analysis and simultaneous consumer evaluations each year. However, due to the low throughput of the assay, sensory characterization is only performed in the final stages of selection prior to release. Alternatively, flavor traits can be incorporated to the breeding program using the breeder’s ‘bite test’, which consists on the individual breeder’s evaluation and scoring of flavors. The major drawback is the fact that the rating generally comes from a single person (n=1), which may not be predictive of the average population overall liking score.

The use of metabolomic profiling as a phenotyping assay can enable accurate characterization of flavor profiles in earlier stages of a breeding program, when more genetic variability is available for selection. For example, the expenditure of $10,000 from a breeding program to run sensory panels in 10 varieties per year, prior to variety release, would allow nearly 200 samples to be phenotyped using metabolomic selection (Figure 6). Metabolic profiling at earlier stages of the breeding program opens up the possibility of identification of superior flavor genotypes that may otherwise have been discarded. Chemical profiling of fruits can capture the genetic potential of the variety as well environmental variability. To reduce this variability and generate a phenotype that better represents the genetic potential of the individual, the breeder can characterize replicates from different environments and/or harvests. Replicates within plots in a single experiment, within harvest dates within a season, or even within environments can be pooled prior to running the instrument, reducing the per sample cost and resulting in an average prediction of the genotypic effect. Furthermore, the use of metabolomic selection would complement a molecular breeding program using genomic selection given that it can better predict sensory attributes using small training sizes derived from sensory panel evaluations. One alternative to combine these approaches would be the application of GS in early stages of the program, for production traits such as yield and adaptability, followed by the application of metabolomic selection in advanced stages that still retain enough genetic variability to select flavorful cultivars (Figure 6). Finally, genes involved in the biosynthesis of inferred volatiles can be targets for further genetic enhancement of fruit flavor. Using the biochemical data along with genomic tools such as genome wide association studies (GWAS) (15), breeding targets can be identified and used to select superior germplasm by marker assisted selection or for gene editing.

**Figure 6:**
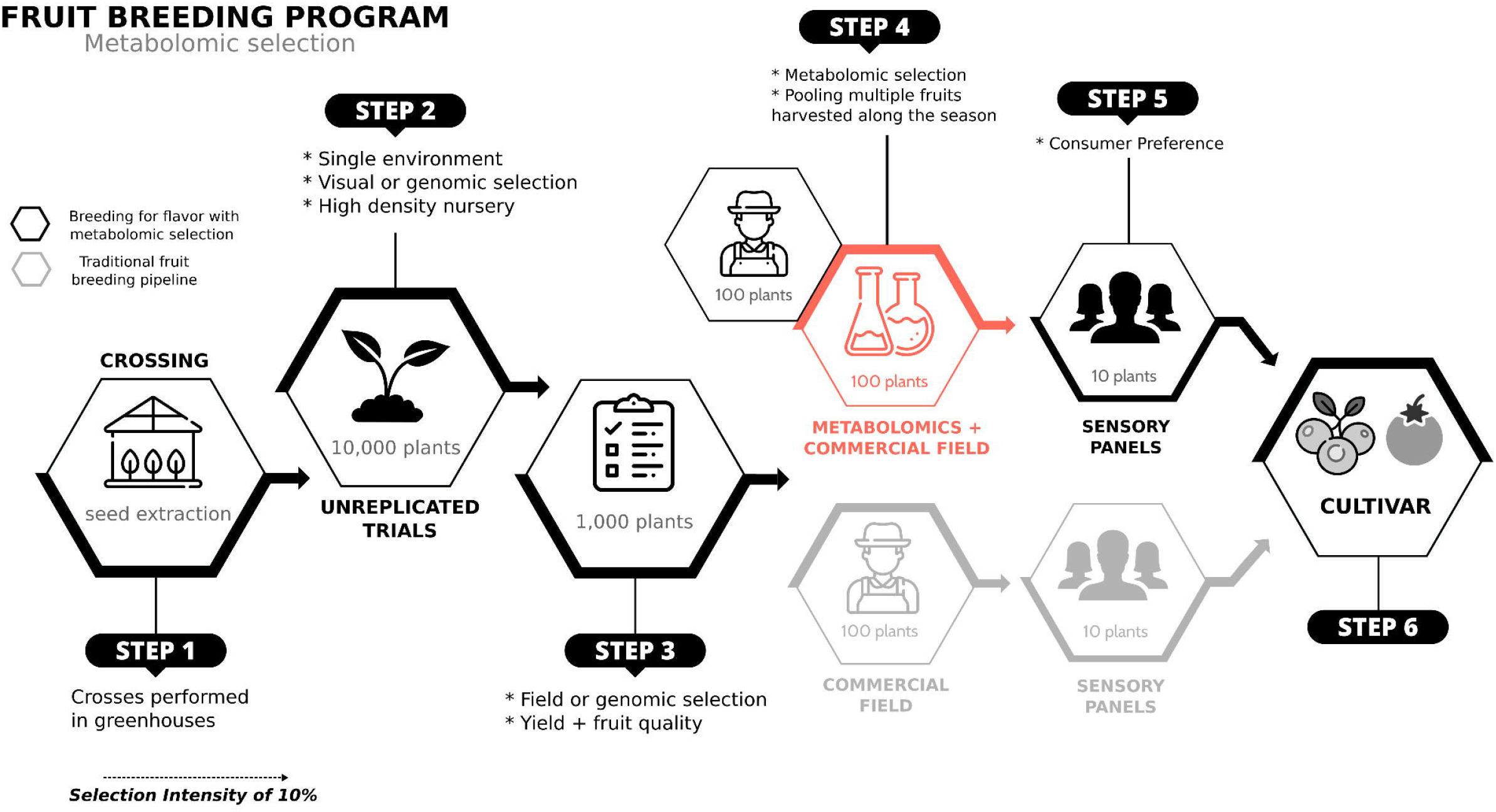
Schematic representation of how the use of metabolomic selection could be applied in earlier stages of a breeding program, when compared to sensory panels.

### 3.3 Machine learning and linear regression models can accurately predict flavor attributes

The use of metabolomics to predict flavor attributes has important implications not only in plant breeding but also in food science and genetics research. Flavor perception is a complex trait as it depends on human preferences. Prediction using metabolomics is challenging due to the correlated nature of the metabolomic predictors since it requires a large number of sensory panels for model calibration. Flavor prediction has been attempted before using linear regression models (26; 27), random forest models (28), and partial least square regression (16), achieving variable levels of prediction accuracies. One of the objectives of our work was to evaluate a range of statistical and machine learning models to determine the best performers for metabolite-based phenotype prediction of flavor quality traits. Importantly, we wanted to access the predictive power of methods known to handle well correlated predictors and thus simultaneously predict the effect of all the variables (metabolites). Identification of the most accurate predictive models provides a simple way to improve phenotyping accuracy with the same dataset available. Here, machine learning models such as gradient boosting machines and XGBoost were the most predictive across all the traits and in both species, whereas multiple linear regression and partial least squares methods were found to be the least predictive. Considering that partial least squares (PLS) is still the standard in food science applications (10), these proposed predictive models show marked improvements with increases relative to PLS ranging from 3.3% for sweetness in blueberry to 44.6% for umami in tomato. These models will increase flavor phenotyping throughput and thereby help to make more informed flavor selections in fruit breeding programs.

The predictive models were used to infer which volatiles contribute to each flavor attribute. For this determination, a Bayesian model (BayesA) and a machine learning model (gradient boosting machines) were chosen. For gradient boosting machines, the importance of each metabolite was quantified by the reduction in accuracy from a model trained on all metabolites to a model trained on all metabolites except for one. This approach allows us to infer which metabolites are important for flavor, but not whether they are important because they are enhancers or suppressors. Each metabolite’s importance in BayesA is measured by the coefficient betas which are either positive or negative depending on whether the metabolite is an important enhancer, or an important repressor.

To better understand the factors affecting flavor perception in each fruit species, we grouped compounds based on their biochemical classification and estimated a proportion of the phenotypic variance associated with each group. Biochemical pathways that were represented by a small number of chemicals were also grouped to minimize the effect of sampling variance in the creation of distance (variance/covariance) matrices (29). As would be expected, sugars (glucose, fructose, soluble solids) were important predictors of sweetness as well as overall liking in both crops, while acids explained a large portion of the sourness variance. By grouping volatiles by their biochemical pathway, we were able to estimate a proportion of the total variance jointly explained by the chemicals within each group. Phenylalanine- and lipid-derived volatiles explained a large fraction of flavor variance in tomato, while lipid-derived, esters, and carotenoid/terpenoid volatiles explained most of the blueberry variance for liking score. Interestingly, ester compounds were shown to be negatively selected in red-fruited tomato as compared to related green-fruited species (30; 31), which potentially explains the lack of contribution to liking score in tomato contrasting to blueberry. Although tomato is botanically a fruit, it is not used as such in most cuisines. Thus, the fruity esters that are so important for flavor in most fruits, do not serve the same function in a tomato.

Multiple regression analysis of volatiles associated with sweetness identified three that contributed to sweetness independently of sugars, but the relative contribution of these volatiles was not determined (4). By grouping volatiles by biochemical pathway and using a linear mixed model, the important role of volatiles in sweetness perception was highlighted. These results also emphasize the need to include volatile detection in breeding programs because by focusing only on sugars and acids during breeding, a large part of the flavor profile is lost (Figure 3). On the other hand, the results suggest that the magnitude of the effect of each individual volatile is much smaller than the individual effect of sugar compounds, highlighting the complexity of breeding for fruit flavor and the challenges to improve flavor using marker-assisted selection. The important contribution of volatiles to overall liking of tomatoes is illustrated by the negative effects of extended refrigeration on volatile contents and consumer preferences (32). Refrigeration substantially reduces volatile contents but not sugars or acids (33).

### 3.4 Conclusions and future directions

In this work we demonstrate the comparison of different algorithms to predict consumer preferences. This information can benefit plant breeding programs to improve flavor perception of new varieties. It is important to note that while we believe that the approach outlined here is generally utile, the specific chemical contributions to overall liking will likely vary with the ethnic and geographic makeup of the consumer panel. While flavor perception is complex due to the subjectivity of the consumers, there are metabolites whose contents can be increased or decreased to improve flavor perception. Future extensions of this could include the modeling of information and parameters for each individual in the panel, such as the inclusion of demographic parameters to predict more nuanced variations in taste preferences. In summary, by creating predictive models for consumer perceptions of flavor we are able to increase the throughput of flavor phenotyping and provide new tools to make more informed, flavorful selections in breeding programs. Through inference, candidate flavor enhancers and suppressors were identified, indicating the possibility of their use as natural food additives in the food industry. Furthermore, genes involved in biosynthesis of these flavor enhancing/suppressing metabolites can now be targets for marker assisted selection or direct engineering of more flavorful fruit varieties.

## 4 Materials and Methods

### 4.1 Data

Prediction analysis was carried in two fruit species: tomato (*Solanum lycopersicum*) and blueberry (*Vaccinium* spp.). For tomato, 68 sugars, acids, and volatiles were analyzed in 147 genotypes grown and evaluated in multiple seasons. A total of 209 samples were used, with 160 samples having been previously evaluated (4) (5). For blueberry, firmness and 55 sugars, acids, and volatiles were analyzed. Sixty-three were grown and evaluated in multiple seasons for a total of 244 samples, of which 164 were evaluated previously (16). Fruit flavor of tomato and blueberry accessions was assessed by consumer sensory panels. Flavor attributes including sweetness, sourness, flavor intensity, and overall liking were rated. Additionally, sensory perceptions of umami were quantified solely for tomato. Overall liking was rated on a scale from −100 to 100 while the remaining attributes were rated from 0 to 100 (3). All data were normalized to a mean of 0 and a variance of 1 for further analyses. Volatiles concentrations were quantified by gas chromatography as described in (17) while sugars, soluble solids, and acids were quantified as described in (34). Sensory analysis was conducted as described in (4) and raw data can be found in Supplemental Data 1 and 2. All blueberry data collection was described in (16). Network analysis was performed using the R package WGCNA (35). Briefly, the pairwise Pearson correlation coefficient between each pair of metabolites was used to construct a weighted metabolite co-expression network. The process assumed an unsigned network and the network was visualized and represented using Cytoscape 3.7.1 (36).

### 4.2 Calculating contributions of metabolites to flavor ratings

To estimate the proportion of variance in flavor ratings that each metabolite group explains, we divided the metabolites identified in tomato and blueberry in six (non-aromatic amino acid derived, Carotenoid/Terpenes, Lipid derived, Phe derived, Sugar and Acids) and seven groups (non-aromatic amino acid derived, Carotenoid/Terpenes, Lipid derived, Ester, Phe derived, Sugar, and Acids), respectively. In tomato, for example, we fit a linear model in which:

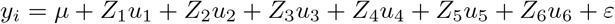

where *y_i_* is the averaged consumer rating for cultivar *i, μ* is the fixed model intercept, *ε* is a normally and independent random effect; **Z** are design matrices for random effects associated to each biochemical group; and **u** are the random terms associated to the chemical groups. For each random term *u* we assumed 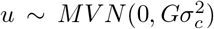, where *G*(.) represents the Gaussian kernel matrices built as the pairwise Euclidean distance between each chemical in a given group. The proportion of the variance explained by a given metabolomic group was determined by 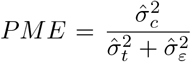; where 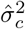 is the variance component estimated for a given metabolomic group; and the denominator is the variance represented by the sum of the variance explained by all other chemical groups 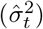 and the residual term 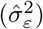. To further represent the contribution of sugar/acids versus volatiles, we also presented it separately by summing the variance components estimated within each group. All analyses were carried out using the `ASReml-R` package (37).

### 4.3 Comparing genomic and metabolomic selection

In order to compare prediction performed via genomic and metabolomic, we organized a group of 70 tomato accessions where whole genome sequencing data, metabolomic evaluation, and sensory data were performed. The genomic data comprised 20 million SNPs, which were mapped to the *Solanum lycopersicum* reference genome as described in (5). We applied additional quality filters and retained only biallelic SNPs with minor alleles frequencies ⩾ 0.1, excluded markers mapped on the chromosome 0, and considered no more than 30% missing data and 20% of heterozygosity rate. Using SNPRelate R package (38) we removed redundant SNPs by pruning markers defined as *r*^2^ ⩾ 0.9 in a 100 kilobase (kb) genome windows. After this step, we retained 79,000 SNPs used in the genomic prediction steps.

Sensory traits were predicted using genomic best linear unbiased prediction (GBLUP) models and compared to metabolomic predictions. The general model for genetic values is *y* = *μ* + *Zg* + *ε*, where **y** is the vector of observed values, *μ* is the fixed model intercept, **Z** is a design matrix that relates the vector **g** of random genetic effects. For genomic prediction, the **g** random effect has null mean and a kernel covariance matrix (**G**) that represents the realized relationship among individuals computed as described by (21). For metabolomic prediction, the kernel kernel was defined in the Euclidean space as described by (39). Residuals were defined as normally and independent distributed. As the number of accessions with metabolomic data is larger than the number of genotyped individuals, we considered two scenarios for the metabolomic data: the *Metabolomic70* represents prediction performances considering the same number of individuals for the genomic and metabolomic prediction (70 individuals) and *MetabolomicAll* represent metabolomic predictions with more individuals (200 individuals). To evaluate the prediction performance of each model, a 10-fold cross-validation was employed, as described in the sequence.

### 4.4 Cross-Validation

To evaluate the predictive performance of each model, a 10-fold cross-validation was employed. In this way, the dataset was randomly split into ten equal groups of varieties. For each of the ten iterations, nine groups of varieties are used to train the model (training set), and one unseen group of varieties is used as a “holdout” group to test the model (test set). During training, a secondary, nested 10-fold cross-validation was used to calibrate the tuning parameters of the machine learning models. The RMSE between predicted and observed flavor ratings in the secondary test set was minimized to obtain the optimal parameter values for the primary model. The trained models were then applied to the metabolite concentrations of the varieties within the primary test set and predicted flavor ratings were obtained. The correlation of the predicted flavor ratings with that of the observed flavor ratings for those varieties is recorded. The average correlation of predicted and observed flavor ratings in the test set is referred to here as the accuracy of the model.

### 4.5 Models

A diverse sample of 18 statistical and machine learning models representing a range of regression, regularization, genomic selection, decision tree, and neural network models were chosen for assessment. These include a linear model and partial least squares as our baseline models; regularization methods such as ridge regression, elastic net, and lasso; kernel methods such as support vector machines, relevant vector machines, and reproducing kernel Hilbert space; neural network models such as a multi-layer perceptron neural network and a Bayesian neural network; decision tree models such as random forest, gradient boosting machines, and XGBoost; and models frequently used in genomic selection such as BayesA, BayesB, and BayesCpi. Each model has their individual strengths, weaknesses, and assumptions. Here we assess which models are most useful for the application of flavor phenotyping by metabolomic selection. All models were implemented in R (40). The Bayesian models were implemented in BGLR (41) and the machine learning models were implemented with caret (42) and a package specific to each model.

#### 4.5.1 Model Descriptions

In this section, we provide a brief overview of the statistical and machine learning methods applied in our case study emphasizing conceptual differences and similarities between approaches. To simplify exposition, the methods have been combined in six classes. For *i* = 1,…, *n*, the model for the *i*-th observation of the response variable *y_i_* corresponding to the vector of features *x_i_* may be typically written as *y_i_* = *f*(*x_i_*) + *ε_i_*, where *f*(.) is the unknown mean function that needs to be learned from the training data, and *ε_i_*, is the *i*-th error. In many instances (particularly, in likelihood-based methods), one often makes parametric assumption about the joint distribution of *ε_i_*’s, e.g., multivariate normal with mean 0 and covariance matrix equal to a multiple of the identity matrix). We assume that the dimension of features is *p*.

#### 4.5.2 Classical (frequentist) multiple linear regression with dimension reduction (LM, PLS

Classical (frequentist) multiple linear regression (MLR) approach assumes that the mean function *f*(*x_u_*) is linear in the covariate coefficients beta, i.e., *f*(*x_i_*) = *x_i_β*. Estimation is typically achieved by ordinary least squares. Dimensionality reduction in the variable selection is achieved by exhaustive enumeration (all subsets when p is small) and by heuristic approaches (such as forward or backward selection) otherwise. Partial least squares (PLS) is a method that is closely related to the principal components regression, with the critical difference that the newly engineered features that are orthonormal linear combination of the original features are created in a supervised fashion, i.e., in explaining the variation in the response vector **Y**, rather than the original design matrix **X**.

#### 4.5.3 Classical (frequentist) penalized/regularized regression

Regularized regression may be viewed as an extension of unpenalized regression that is particularly appropriate when the number of covariates is large. Specifically, the objective function for minimization in order to estimate the model parameters is the SSE plus a regularization penalty on the beta coefficients, which has an effect of shrinking the coefficient toward zero. Specific examples are ridge regression, lasso regression and elastic net. One attractive operating characteristic of the lasso regression is that it often results in the exact shrinkage of a subset of the coefficients to zero, which achieves variable selection.

#### 4.5.4 Bayesian regression for Genomic Selection

The methods presented in this subsection may be viewed as the Bayesian extensions of the penalized regression approaches presented earlier. In the Bayesian approaches, information about parameters and latent variables (such as random effects) is expressed using prior densities on those parameters. Inference is typically performed numerically using Bayesian computation machinery via Markov Chain Monte Carlo sampling. These inference schemes require the specification of the joint density of the data given the parameters and latent variables, as well as the priors on the latent variables and the model parameters (typically, on the log-scale for reasons of numerical stability). It is notable that the ridge regression method is equivalent to placing a multivariate normal prior on beta, while the lasso penalty is equivalent to the Laplace (double exponential) prior on beta. To replicate the model/variable selection capability of the frequentist lasso, one often considers spike-and-slab priors on the vector of coefficients beta; these are a mixture of a multivariate normal (or Student’s t) or Laplace prior on beta (the slab part) and a point mass at zero (the spike part). These choices are implemented in the BGLR package as Bayesian lasso and ridge regression, and Bayes A, B and C. We additionally evaluate the BGLR implementation of the Reproducing Kernel Hilbert Space model which is effectively a Gaussian process model with a covariance function induced by the kernel function *K*(.,.).

#### 4.5.5 Ensembles of simple regression trees (random forest, gradient boosting)

Regression tree is a decision tree algorithm aiming at partitioning the predictor space as a union of disjoint hyper-rectangular regions, with the prediction on the *j*-th region being the sample mean of the responses from the training data whose features fall into the *j*-th region. While individual regression trees are often interpretable, predictions from single trees are generally not competitive with other machine learning methods. This deficiency is often overcome with ensembles of trees, where hundreds or thousands of simple trees (with low depth of interactions/small number of splits) are fitted to the training data, and a consensus prediction from the entire ensemble is used. Random forests fit the collection of trees to bootstrap resamples of the training data subject to constraints on the choice of splitting variables for each split, in order to “decorrelate” the trees in the ensemble. Gradient-boosted regression trees proceed iteratively, by “underfitting” the residual from the fit in the previous iteration using a shrunken version of a newly fitted tree; the rate of underfitting is controlled by a separate tuning parameter known as the “learning rate”. Unlike single-tree models, trees in the ensemble need not be deep, and in many instances one or two splits per tree outperform deeper models.

#### 4.5.6 Separating hyperplanes/support vector machines

In the simplest case, separating hyperplane methods aim at constructing a linear decision boundary to separate two classes in the binary classification problem. Maximum margin classifiers allow one to extend the formulation to allow one, “softly” and in a more stable fashion, to separate the classes when a limited amount of constraint violation is allowed. Basis expansions and kernel methods such as the classical support vector machines (with nonlinear kernels, e.g., radial basis function with polynomial and Gaussian bases) allow more flexible separation of the classes thanks to the nonlinear decision boundaries that they provide. Further extensions to address the regression (as opposed to the classification) problem have been implemented in the R library kernlab (43); here, the mean function of the quantitative response *y_i_* is modeled as a linear combination of basis functions expressed using the kernel, in a fashion similar to the splines. Relevance vector machine further extends this specification using a Bayesian framework, allowing the basis function coefficients and hyperparameters to be calibrated in a Bayesian fashion.

## 6 Author Contributions

MFRRJ designed the study. MFRRJ, PRM and HK coordinated the study. DT and HK coordinated the genomic and metabolomic data collection for tomato. PRM and LFVF organized the metabolomic historical data collected by the blueberry breeding program. VC and LFVF performed the data analyses. CS provided expertise on the sensory analysis performed in blueberry and tomato. NB provided statistical expertise in machine learning methods. VC, LFVF, HK, DT, and MFRRJ contributed with data interpretation and wrote the paper. All authors read and approved the final version of the manuscript for publication. VC and LFVF contributed equally to this work.

## 7 Acknowledgments

The authors thank previous students that, as part of the tomato and blueberry UF breeding program, supported the panel data collection and volatile extraction. This work was supported by the UF royalty fund generated by the licensing of blueberry cultivars, by the National Science Foundation (IOS 1844237 and IOS 1855585 to HK and DT, and IOS 1564366 to DT), and the National Institute of Food and Agriculture (SCRI 2018-51181-28419 to MFRR).

