## Supplementary material for "Metabolomic Selection for Enhanced Fruit Flavor": Figure S1

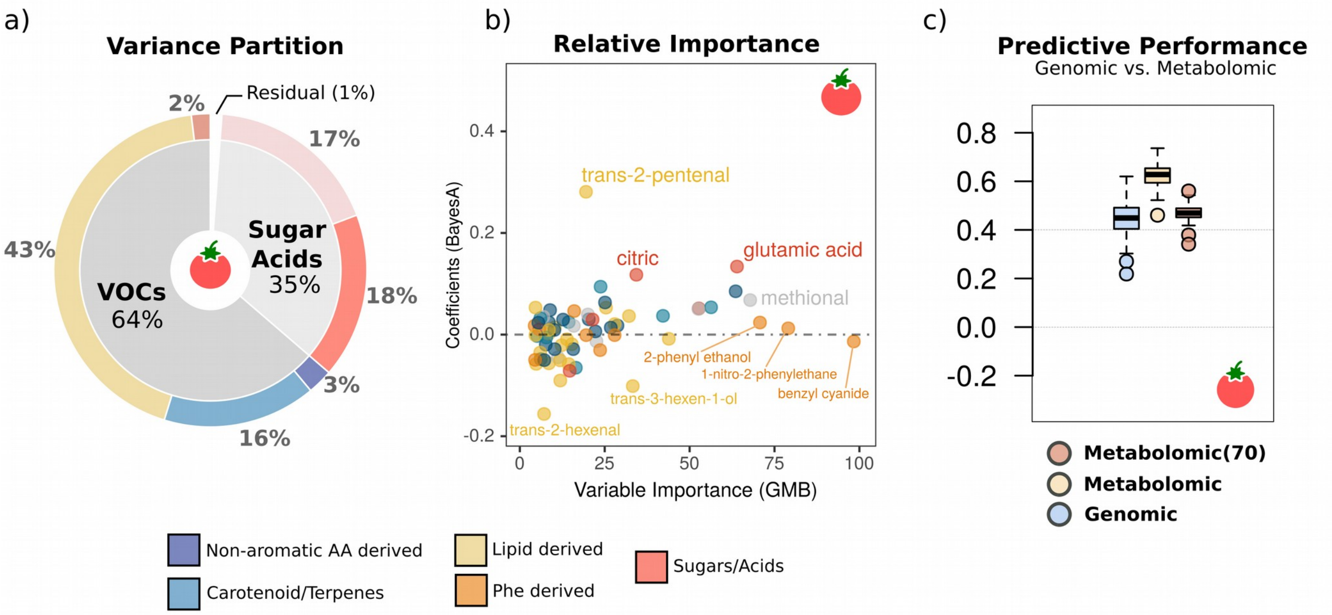


Figure S1: a) Variation in sensory umami ratings explained by sugars/acids and volatiles overall, and by groups of metabolites of known biochemical classification in tomato; b) Relative importance estimated with Gradient Boost Machine (x-axis) and effect level estimated with BayesA (y-axis) of each metabolite in predicting umami ratings in tomato. Colors indicate the type of metabolite; c) Accuracy of predicting umami sensory trait for tomato using 70 individuals with genomic (Genomic 70) and metabolomic (Metabolomic 70) data, as well as all 209 individuals with metabolomic data (Metabolomic All)
